# Local overfishing patterns have regional effects on health of coral, and economic transitions can promote its recovery

**DOI:** 10.1101/2021.08.15.456395

**Authors:** Russell Milne, Chris Bauch, Madhur Anand

**Affiliations:** Department of Applied Mathematics, University of Waterloo; School of Environmental Sciences, University of Guelph

**Author notes:** Corresponding author Email address (Russell Milne).

**Keywords:** Coral reefs, overfishing, transient dynamics, human-environment model, habitat fragmentation

## Abstract

Overfishing has the potential to severely disrupt coral reef ecosystems worldwide, while harvesting at more sustainable levels instead can boost fish yield without damaging reefs. The dispersal abilities of reef species mean that coral reefs form highly connected environments, and the viability of reef fish populations depends on spatially explicit processes such as the spillover effect and unauthorized harvesting inside marine protected areas. However, much of the literature on coral conservation and management has only examined overfishing on a local scale, without considering how different spatial patterns of fishing levels can affect reef health both locally and regionally. Here, we simulate a coupled human-environment model to determine how coral and herbivorous reef fish respond to overfishing across multiple spatial scales. We find that coral and reef fish react in opposite ways to habitat fragmentation driven by overfishing, and that a potential spillover effect from marine protected areas into overfished patches helps coral populations far less than it does reef fish. We also show that ongoing economic transitions from fishing to tourism have the potential to revive fish and coral populations over a relatively short timescale, and that large-scale reef recovery is possible even if these transitions only occur locally. Our results show the importance of considering spatial dynamics in marine conservation efforts, and demonstrate the ability of economic factors to cause regime shifts in human-environment systems.

## Introduction

Coral reefs are home to very high levels of biodiversity [1, 2, 3], and provide vital services to humans such as harvesting of reef fish and ecotourism [4]. Overfishing has long been known to be a major stressor of reefs [5, 6, 7], due to its ability to shift areas from a coral-dominated to a macroalgae-dominated state [6, 8]. Such shifts, when considered on a regional scale, can disrupt connectivity on a reef and give rise to fragmented rather than connected habitats. This has been shown to alter the composition of species present [9, 10], although the overall effects of fragmentation are ambiguous [11]. Additionally, as the economies of reefside communities transition from being based on fishing to tourism, areas that were previously overfished may see a regime shift in the opposite direction, back to coral dominance. However, the speed of such a shift, as well as whether one can happen regionally due to local-scale economic transitions, is yet to be seen. Here, we use a spatially explicit coral reef model using a coupled human-environment framework to investigate these multi-scale processes, and to determine their implications for the future viability of both coral reefs and the communities that depend on them.

Overfishing of reef fish has been cited as one of the most prominent threats to the livelihood of coral reefs (e.g. [5, 6, 7]). This is due to the fact that many commercially valuable species of reef fish, and especially parrotfish, are predators of macroalgae [12, 13], which can overgrow coral and outcompete it for available space. Fishing pressure on heavily-harvested coral reefs in the Pacific has been estimated at or above 50 percent of organisms from many different commercially viable species per year [14, 15]. Many of the species surveyed were being fished above levels predicted to be sustainable, including half of all parrotfish species in Hawaii [14], and harvesting rates in general were often far above the rates associated with a shift to a macroalgae-dominated state according to past modelling results [8, 16]. Further complicating matters is the fact that other coral reef stressors, such as nutrient loading, have interacting effects with overfishing that increase the propensity of an overfished system for a regime shift even further [17, 18].

While overfishing is known to have deleterious effects on coral reefs, including causing regime shifts, fishing can safely be performed at lower rates without these risks. Harvesting rates associated with small-scale subsistence fishing, which have previously been estimated at one tenth of commercial rates [19], have been found to be between one seventh and one third of the estimated upper limits for sustainability of coral populations [20]. Many communities situated adjacent to coral reefs are in the process of transitioning from economies based around commercial fishing to those more heavily based around tourism, including those in the Pacific [21, 22] and the Caribbean [21, 23]. After this transition, fishing operations would typically be on a smaller scale; the wide gap between commercial and subsistence fishing rates suggests the possibility that these economic transitions could drive regime shifts. In particular, this raises the question of how quickly a reef that has previously been under very high fishing pressure can recover following such an economic transition. In addition to this, as commercial fishing is an important industry both in terms of how much revenue it generates [4, 24, 25] and how many people depend on it for food [6, 24], it is necessary to balance the needs of the fishing and tourism industries as well as the coral reef itself. Ideally, a reefside community should have a healthy reef as well as sustainable fishing and tourism industries; therefore, finding conditions for the coexistence of these is imperative.

In addition to featuring a wide array of trophic interactions such as the linkages that cause coral to be harmed by overfishing, coral reef ecosystems are also very complex spatially. Part of this is due to their sheer size. The Great Barrier Reef is the largest marine protected area in the world [26], and the Caribbean Sea similarly features a large network of reefs offshore of various islands. Reefs within a given region are also incredibly heterogeneous in their internal composition, with sites dominated by coral, macroalgae, and algal turf all being present [27]. Similarly, different reefs, and different areas of a reef, are highly connected due to the dispersal of coral larvae [28, 29], fish [30, 31, 32] and nutrients, and these dispersal processes themselves have different effects across different spatial scales [29]. Therefore, damaging one part of a reef also should have farther-reaching effects on areas that it is connected to. However, most modelling of overfishing and other coral reef stressors has been done strictly at local scales (see review in [33]). Because of this, an increased focus on multi-scale effects of overfishing has the potential to uncover many new insights.

Owing to coral reefs’ size and complexity, concepts pertaining to nonlocal processes are often seen in field and theoretical literature related to reefs. For instance, previous models considering connectivity between coral reef habitats implicitly [34] and explicitly [35] have emphasized the importance of the spillover effect, where dispersal of coral larvae or fish from relatively undisturbed reefs into adjacent fished areas can help counteract the degradation caused by overfishing. A potential counteracting effect is large-scale and commercial harvesting in areas that are nominally protected, as seen with many species associated with coral reefs [36, 37, 38]. Fishing boats routinely travel sizable distances away from their home ports [39, 40], and outside fishers often employ overly damaging fishing techniques against marine protected area (MPA) regulations [36]. Hence, an MPA without enforced boundaries is liable to have substantial fishing pressure from adjacent areas outside it. In light of this, it is important to understand how processes such as nonlocal fishing pressure and the spillover effect can interact to affect reef health over broader spatial scales.

Echoing the spatial heterogeneity present in coral reef ecosystems, the debate over the best conservation strategy for coral is also spatial in nature. Habitat connectivity has been cited as important for the design of marine protected areas (MPAs) on coral reefs [41, 42, 28], as it also has with other types of marine ecosystems [43], and its prominence in the literature has been steadily increasing over time [44]. However, the debate over the importance of connectivity is not closed, as it rests on the distances that the species being protected can disperse. One recent paper concluded that the dispersal abilities of Caribbean reef fish are insufficient to traverse the gaps between current MPAs [32], whereas other work has found that marine species disperse over such great distances that the importance of connectivity in designing MPAs is minimal [45]. Because of these discrepancies in the literature, and given the importance of establishing sound conservation strategies for coral reefs, more research on the optimal configuration of MPAs is needed.

Analogous to the debate over connectivity of MPAs is that over the relative threats posed by habitat loss and habitat fragmentation. Again, this is underpinned by the dispersal abilities of the species that would be protected. Although habitat fragmentation is a great concern in terrestrial ecosystems, marine species generally have greater dispersal ability and are therefore affected less by it. Fragmentation has been shown to have highly variable effects on the functioning of coral reefs and other marine ecosystems [11], including on abundance and biodiversity of reef fish [10], and the effects of habitat fragmentation via degradation due to overfishing may also be countered by mechanisms such as the spillover effect. Recent calls have been made for more research on the variety of responses that marine communities have to fragmentation, especially research that integrates dynamics over both local and regional scales [11]. Hence, it is necessary to build a robust, multi-scale theory around how important connectivity and fragmentation are for the viability of the many species that inhabit coral reefs.

In this paper, we use a coupled human-environment model to determine the effects of overfishing on coral reefs across both local and regional scales, and provide policy solutions for managing overfished reefs. We identify how economic transitions can lead to regime shifts from macroalgae to coral dominance, and show that these transitions, when occurring locally, can promote both healthy reefs and a sustainable economy with fishing and tourism both being viable. We test the ability of coastal communities to stop coral decline via temporarily subsidizing the tourism industry, and find that such short-term subsidies can drive long-term coral recovery. We contrast the spatial effects of fish and coral dispersal with those of nonlocal harvesting inside MPAs, and show the importance of strict enforcement of MPA boundaries. We also determine that coral and herbivorous fish have very different responses to habitat fragmentation, with the implication that MPA design needs to take into account divergent needs of multiple species.

## Methods

### Model formulation

To simulate the dynamics of a coral reef, we adapted a dynamical system model of Spiecker et al. [35] featuring herbivorous fish, coral, macroalgae, nutrients and detritus. We chose this model because it is mechanistic and based around recruitment and mortality rates (as opposed to state transition models, e.g. [46]), which is important as the regional-scale dynamics we investigate strongly involve processes like the production and dispersal of coral larvae. In order to be able to capture the dynamics in many different areas of a reef habitat, we simulated a linear network of patches along a coastline, interconnected via dispersal of the model’s components. This is similar to the integro-differential equation approach used in [35], although we assumed a patchy rather than continuous landscape in order to capture the effects of overfishing within a certain area rather than at a single point. In order to perform an in-depth examination of overfishing, especially as it relates to economic transitions between fishing-based and tourism-based economies, we added a novel differential equation to the model representing the proportion of economic activity in each patch related to tourism (rather than fishing), with change over time driven by the relative economic utility gained from these strategies. We also introduced a dynamic fish harvesting rate that depends on economic strategies in each patch and incorporates harvesting by fishing boats outside their local patch. Our full human-environment model is below:

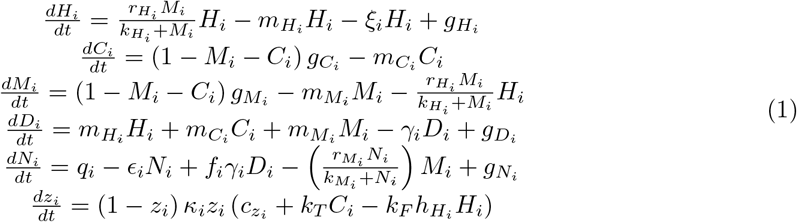

The dispersal functions in the model are below. For coral and macroalgae, this is part of their growth rate, whereas for the other components this is mathematically equivalent to passive dispersal.

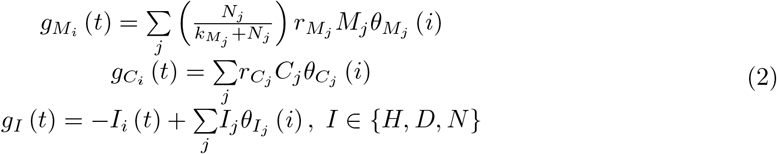

The dynamic harvesting rate is below:

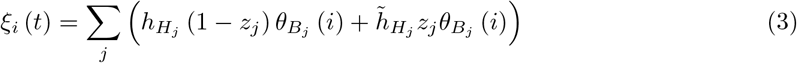

Within the model, there are five biological components. Each one represents a certain functional group or abiotic factor rather than focusing on individual species, an approach also used in e.g. [46, 47]. These are herbivorous fish *H*, coral *C*, macroalgae *M*, detritus *D* and nutrients *N*. Coral and macroalgae compete for space offshore, and therefore their total abundance is restricted in the model, i.e. *M* + *C ≤* 1. Any space not colonized by coral or macroalgae is assumed to be covered by algal turf or bare rock. The herbivorous fish population has been normalized to be on the same scale as coral and macroalgae, and hence is expressed in terms of its density over an arbitrary area. (Scaling constants with units of area^*−*1^ are therefore omitted due to being equal to 1).

In the model, herbivorous fish are assumed to eat macroalgae using a Holling Type II functional response, represented as a mathematically equivalent Hill function with maximum growth rate *r*_*H*_ and half-saturation constant *k*_*H*_. The fish reproduce at the rate at which they eat macroalgae, die of natural causes at a rate *m*_*H*_, and are harvested at a variable rate (detailed below). Coral and macroalgae reproduce via the dispersal of larvae and propagules [34], so their growth rate is nonlocal. Coral larvae are created in each patch at a rate *r*_*C*_. This is constant in [35], but we took it to vary temporally, since coral reproduction events happen at specific times during the year [48] and to allow for a mechanism for macroalgae to overgrow coral. Macroalgae propagules are created in each patch at a rate depending on the available nutrients, which is governed by a saturation function with maximum growth rate *r*_*M*_ and half-saturation constant *k*_*M*_. At each time step, the new coral larvae and macroalgae propagules are distributed among patches according to their distances from whichever patch the larvae and propagules originated in. Dispersal of new larvae and propagules out of each patch is governed by a Gaussian dispersal kernel centred on that patch that has been discretized (see e.g. [49]), and the intrinsic growth rate for coral or macroalgae in one patch is the sum of the larvae or propagules created anywhere that disperse into that patch. This rate is scaled down by the factor (1 *− M − C*) for both, to represent their shared carrying capacity. Coral die of natural causes at a rate *m*_*C*_, while macroalgae die of natural causes at a rate *m*_*M*_ and are eaten by fish as detailed above. Detritus is formed by organisms that die of natural causes at one-to-one rates, and decays into nutrients at a rate *g*_*D*_. Nutrients are formed from detritus at the same rate, scaled by a conversion constant *f*_*N*_, and are uptaken by macroalgae as mentioned above. Nutrients also enter the system via inorganic processes (e.g. river outflows) at a rate *q*_*N*_ and leave it (e.g. by ocean currents) at a rate *e*_*N*_. These processes can be seen in a schematic of the local dynamics of the model (Fig. 1).

**Figure 1:**
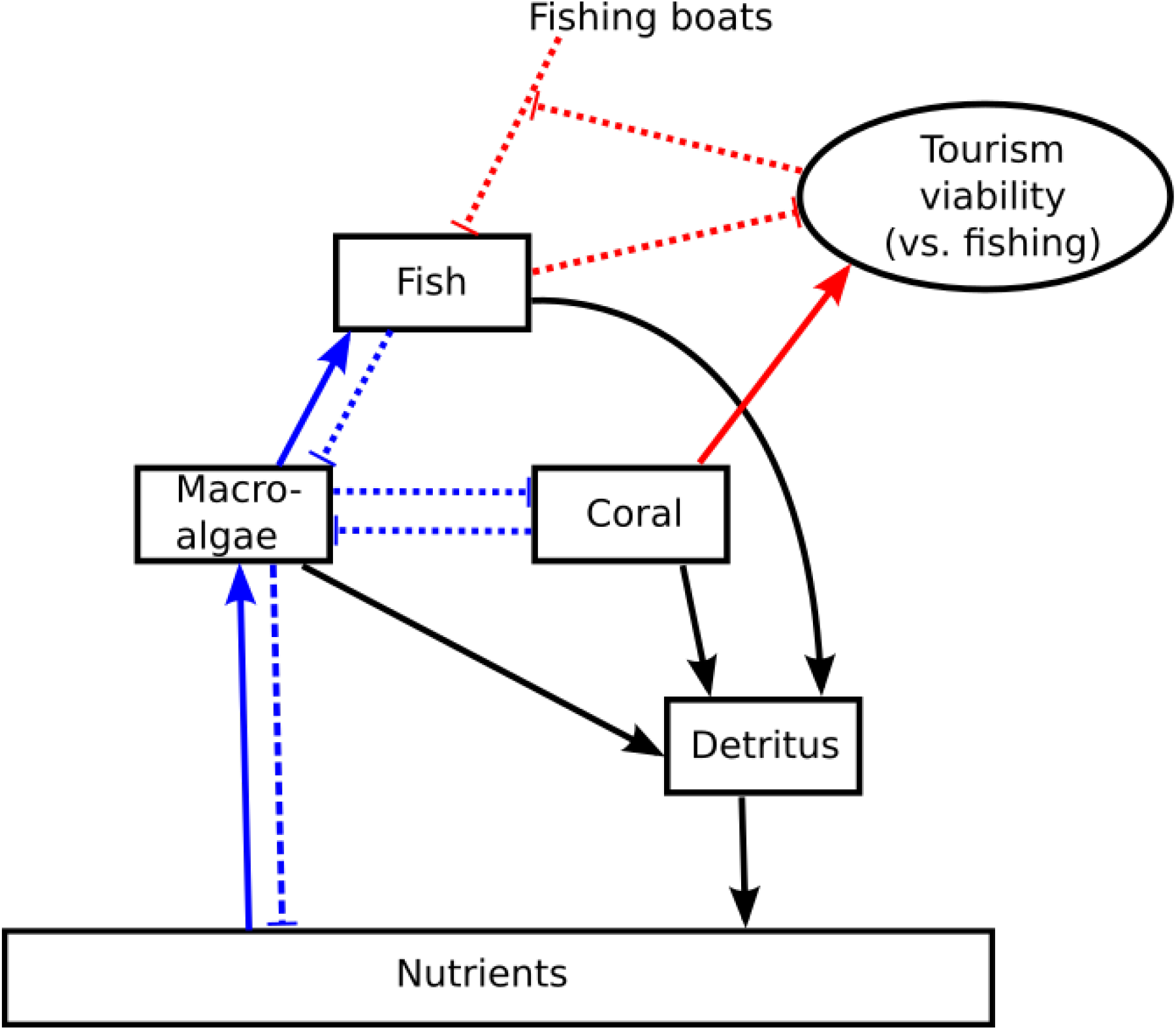
Schematic showing local interactions between model components. Red lines represent economic interactions, blue ones represent trophic and competitive ones, and black ones represent cycling of materials.

In addition to their local dynamics, fish, detritus and nutrients are assumed to undergo passive dispersal between patches. The dispersal rates for these are governed by Gaussian dispersal kernels in the same way as coral and macroalgae reproduction are. Adult coral and macroalgae are assumed not to move. We chose baseline values of the standard deviations of each kernel to be 1. This represents relatively large ability for fish, coral larvae and macroalgae propagules to disperse outside of their home patches. The standard deviations for detritus and nutrients are the same as those for fish, coral larvae and macroalgae propagules since dispersal by the latter groups is dependent on physical factors such as tides and ocean currents, which also drive dispersal by the former groups. A Gaussian dispersal kernel is also used to quantify the amount of time that fishing boats from any given patch spend in each patch in the system (including their own), and hence the nonlocal fishing pressure in each patch; this is denoted *σ*_*B*_.

We complemented the biological components of the model by adding a state variable *z* encompassing local economic strategies, in a format similar to that used previously for quantifying support for conservation [50]. Specifically, we considered two economic strategies, namely fishing and ecotourism, and let *z*_*i*_ be the proportion of economic agents in patch *i* engaging in ecotourism. In our model, *z* changes according to the relative utility of both strategies. *z* increases when a large amount of coral is present (and hence ecotourism is more profitable), and it decreases when large quantities of fish are available to be harvested (measured by the quantity 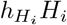). We use the parameters *k*_*T*_ and *k*_*F*_ to scale how strongly support levels for tourism and fishing, respectively, depend on the underlying biological conditions, and we define *c*_*z*_ as the degree to which one strategy is more profitable than the other due to external factors. Additionally, *z* changes due to social pressure using the replicator dynamics found in [50], under the assumption that economic agents in a patch will be more likely to use a particular strategy if their neighbours are also using (and profiting from) it. We take *κ* to be the base rate at which economic agents can switch their strategies. *z* is coupled back into the model via the dynamic fishing rate, as only economic agents engaging in fishing are assumed to fish at the higher commercial rate. This can be seen in the local schematic (Fig. 1).

### Model parametrization

To obtain the kinetic rates for herbivorous fish growth, we surveyed the doubling times of all parrotfish species in FishBase [51], the same method as that used in previous modelling papers (e.g. [16, 50]). We took *r*_*H*_ = 0.7 yr^*−*1^ and *k*_*H*_ = 0.5, since the majority of species were categorized as having doubling times less than 15 months and nearly all of the rest were in the category of having doubling times of 1.4 to 4.4 years. Our values produce a reproduction rate similar to the linear rates used in previous work [16, 50] when macroalgae cover is at its maximum of *M* = 1. Coral reproduction rate for different species has been estimated at annual doubling [52], an average of 5.7 larvae per colony per year [53], and ten eggs per polyp in a yearly spawning session [54]. We took *r*_*C*_ = 5 yr^*−*1^, a value in the middle of this range. Macroalgae are known to grow very quickly, and tenfold yearly growth under optimal conditions has been reported [55]. Spiecker *et al*. deemed a value of 15 for macroalgae growth rate to be biologically plausible, and their sensitivity analysis found most state variables to be minimally responsive to changes in it, so we took *r*_*M*_ to be the slightly lesser value of 12 yr^*−*1^. We kept the value of *m*_*C*_ = 0.44 yr^*−*1^ from previous studies [50], and used the low natural mortality rate of 0.1 yr^*−*1^ for herbivorous fish and macroalgae.

When considering nutrient dynamics on coral reefs, we looked specifically at nitrogen. This was done because macroalgae and other primary producers on pristine coral reefs have shown N-limitation, but those closer to developed areas are often saturated with nitrogen due to anthropogenic input and therefore are P-limited instead [56, 57, 58]. Coral reefs have high rates of nutrient exchange with the surrounding oceanic water [59] and short residence times [60], so we took *E* to be a high rate of 0.6. Nutrient input into coral reefs and other marine ecosystems has been estimated as on the order of 100 to 1000 kg N km^*−*2^ yr^*−*1^ in most areas [61, 26], with higher values for areas of dense human settlement, and nitrogen concentration of water entering wetlands adjacent to the Great Barrier Reef has been measured at 200 *µ*g N L^*−*1^ under flood conditions [62]. We therefore considered values of *q* ranging from 20 to 120 kmol N yr^*−*1^, representing the total amount of nitrogen exported into a patch of approximately 1 km^2^. We took *γ* to be 1 yr^*−*1^ under the assumption that all detritus would decompose within a year [63, 64], and used a value of 20 kmol N for *f* as nutrient input from detritus decomposition was expected to be an order of magnitude less than input from external loading [65]. We used a half-saturation constant for nutrient uptake by macroalgae (*k*_*M*_) of 80 kmol N yr^*−*1^, close to the median of the values reported in previous studies [66, 67] after adjusting units to make *k*_*M*_ on the same scale as *N*. This choice also meant that nitrogen availability was close to saturation at the upper ranges of *q* that we tested, as expected.

We used a baseline of 0.5 for the commercial fishing rate *h*_*H*_; this value has been used in prior human-environment modelling work on coral reefs [50] and is consistent with available data for reef fish harvesting [14, 15]. We took the subsistence fishing rate 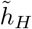 to be 0.05, one tenth of the baseline commercial rate [19]. Unless we were simulating economic transitions or the effects of tourism subsidies (see below), we took *c*_*z*_ to be zero, indicating no external economic pressure in favour of fishing or tourism. We fit the other social parameters (*κ, k*_*T*_, *k*_*F*_) by simulating the system for different orders of magnitude of these parameters, and choosing values for which *z* converged to equilibrium at 0 or 1 after a plausible length of time following a shock. We ultimately took *κ* = *k*_*T*_ = *k*_*F*_ = 1.

### Numerical methods

In order to investigate local dynamics and check when regime shifts are expected to take place, we simulated a one-patch version of the model while varying harvesting rate (*h*_*H*_) and nutrient loading rate (*q*). This allowed us to determine how the ability of overfishing to push coral reefs into a macroalgae-dominant regime is mediated by nutrient loading. For each run of the model, we determined the post-transient average values of coral and macroalgae cover, as well as herbivorous fish abundance. We defined different regimes as discrete regions of parameter space (harvesting rate vs. nutrient loading rate) with qualitatively similar dynamics.

To evaluate whether overfishing-driven habitat loss or fragmentation is more detrimental to coral and herbivorous fish, we simulated a network of 25 patches in which a fixed number of patches were heavily overfished (*h*_*H*_ = 0.8) and the rest were fished at subsistence levels (*h*_*H*_ = 0.05). We varied both the number of overfished patches (to test the effects of habitat loss) and their configuration (to test the effects of habitat fragmentation). Configurations that we used included one where all overfished patches formed a contiguous area in the middle of the simulated landscape, with large contiguous areas of non-overfished patches on either side, and several where overfished and non-overfished patches alternated in a repeating pattern. Each of these patterns involved taking specifying a certain number of patches to be overfished and taking the rest to be non-overfished, and spacing groups of overfished patches evenly throughout the system (where each group consisted of a fixed number of patches). For each run of the model, we took the average post-transient values of coral cover and herbivorous fish abundance across the landscape as a whole, in overfished patches and in non-overfished patches. This allowed us to easily separate the local and regional effects in each scenario. We also took different values of *σ*_*B*_ to control for the effects of nonlocal harvesting, using values of 0.25 (for a system in which fishing is almost entirely done locally) and 1 (for a system in which substantial amounts of harvesting takes place outside of fishing boats’ local patches).

To determine the relative effects of the spillover effect and fishing across MPA boundaries, we determined the average equilibrium values of herbivorous fish and coral in a 25-patch system while varying their dispersal abilities (*σ*_*H*_ and *σ*_*C*_) and the amount of time fishing boats spend locally (represented by the mean value of the discretized Gaussian distribution generated by *σ*_*B*_). In the simulated system, approximately half of the patches (13 of 25) were overfished (*h*_*H*_ = 0.5) and the rest were fished at subsistence rates. The overfished patches were either located in a contiguous stretch in the middle of the simulated area (the “contiguous case”) or alternating one-to-one with non-overfished patches (the “fragmented case”).

To determine the long-term effects of economic transitions between a fishing-based economy and a tourism-based one, as well as check conditions for the coexistence of fishing and tourism, we ran simulations that treated *c*_*z*_ as a time-dependent function, rather than its static baseline value of zero. We ran different simulations to represent long-term economic trends and temporary subsidization of the tourism industry. For long-term trends, we made *c*_*z*_ increase linearly from 0 to a maximum value of 5 over a span of five years. For short-term subsidization, *c*_*z*_ was initialized at a positive constant value (taken to be integer values from 1 to 5), held there until a time 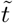 (taken to be integer values from 1 to 15), and then reset to 0. As above, we used a 25-patch system. In the long-term trend scenario, we altered *c*_*z*_ in a varying number of connected patches (1 to 25) in the middle of the simulated area to test the effects of both local and regional economic shifts.

In both long-term and short-term scenarios, we initialized the system using initial conditions representative of the macroalgae-only regime, and determined the amount of time taken before the overfished patches shifted back to a coral-dominated state (defined as over 50 percent coral cover) and healthy fish population levels (fish density of 1). These values were chosen to represent typical average coral and fish levels in the coral-dominated regime (see Results section), and to be high enough that the systemwide average attaining these levels would indicate a regional-scale recovery. We also checked the long-term average coral cover in these systems, to test whether recovery was temporary or permanent. The initial conditions that we used in overfished patches were 90-99 percent macroalgae cover with the rest of the seabed covered by coral, fish density equal to the amount of coral cover, fishing being 99 percent of the economic activity, and detritus and nutrients being at their average steady-state levels reached under these conditions. We also took *h*_*H*_ as the constant value of 0.5 in each patch to preclude the possibility of natural recovery.

## Results

### Local dynamics and regime shifts

We found three distinct regimes that the system’s local dynamics can take (Fig. 2). The first of these featured cyclical dynamics, with coral dominant most of the time and macroalgae always present. In this regime, tourism eventually composed all economic activity (Fig. 3), as *z* rose and fell depending on the relative abundances of coral and fish but was always higher at the end of a cycle than at its beginning. Due to the lack of fishing pressure, the herbivorous fish and macroalgae populations followed oscillatory boom-bust patterns similar to those found in the Rosenzweig-MacArthur model. The second regime featured stable, nonzero levels of coral, macroalgae and herbivorous fish. Here, macroalgae was dominant over coral, with coral cover of the seabed typically above 10 percent but below 30 percent. Economic activity converged to a state where only fishing was viable, although very long transients were possible depending on the social parameters (Fig. 3). However, fish populations were higher in this regime than they were in the cyclical coral-dominant regime. The third regime was characterized by local extinction of both coral and herbivorous fish, with macroalgae taking up all available space on the seabed. Economic activity tended towards the all-fishing equilibrium while there were still fish available to catch. However, after a certain point in time, changes in economic behaviour became minimal as coral and fish populations were both roughly zero and no economic utility could be gained from either of them.

**Figure 2:**
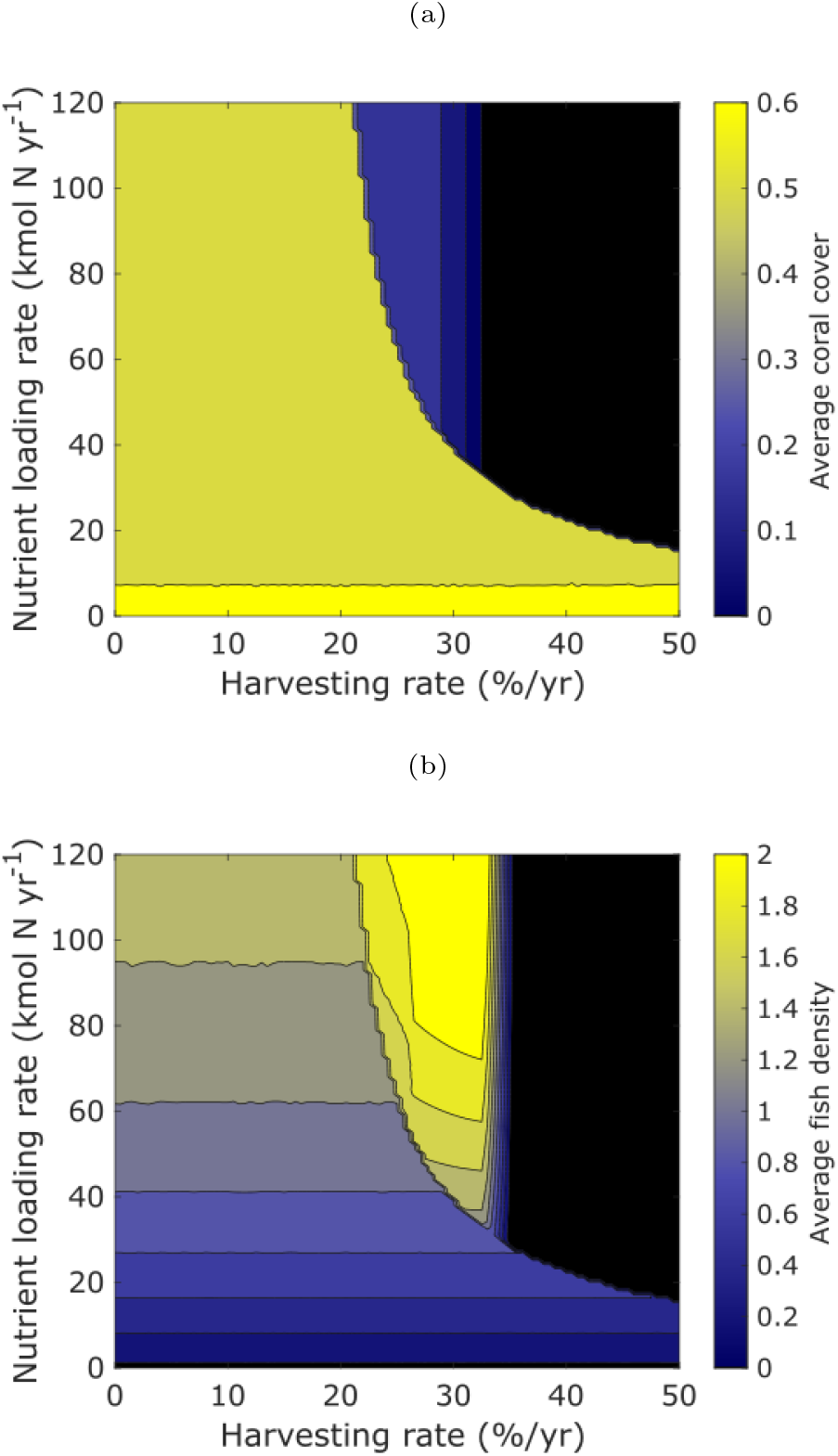
Levels of coral cover (Fig. 2a) and herbivorous fish density (Fig. 2b) in one patch as a function of harvesting rate and nutrient loading rate, showing three distinct regimes. Values taken are the equilibrium value or the average over one limit cycle.

**Figure 3:**
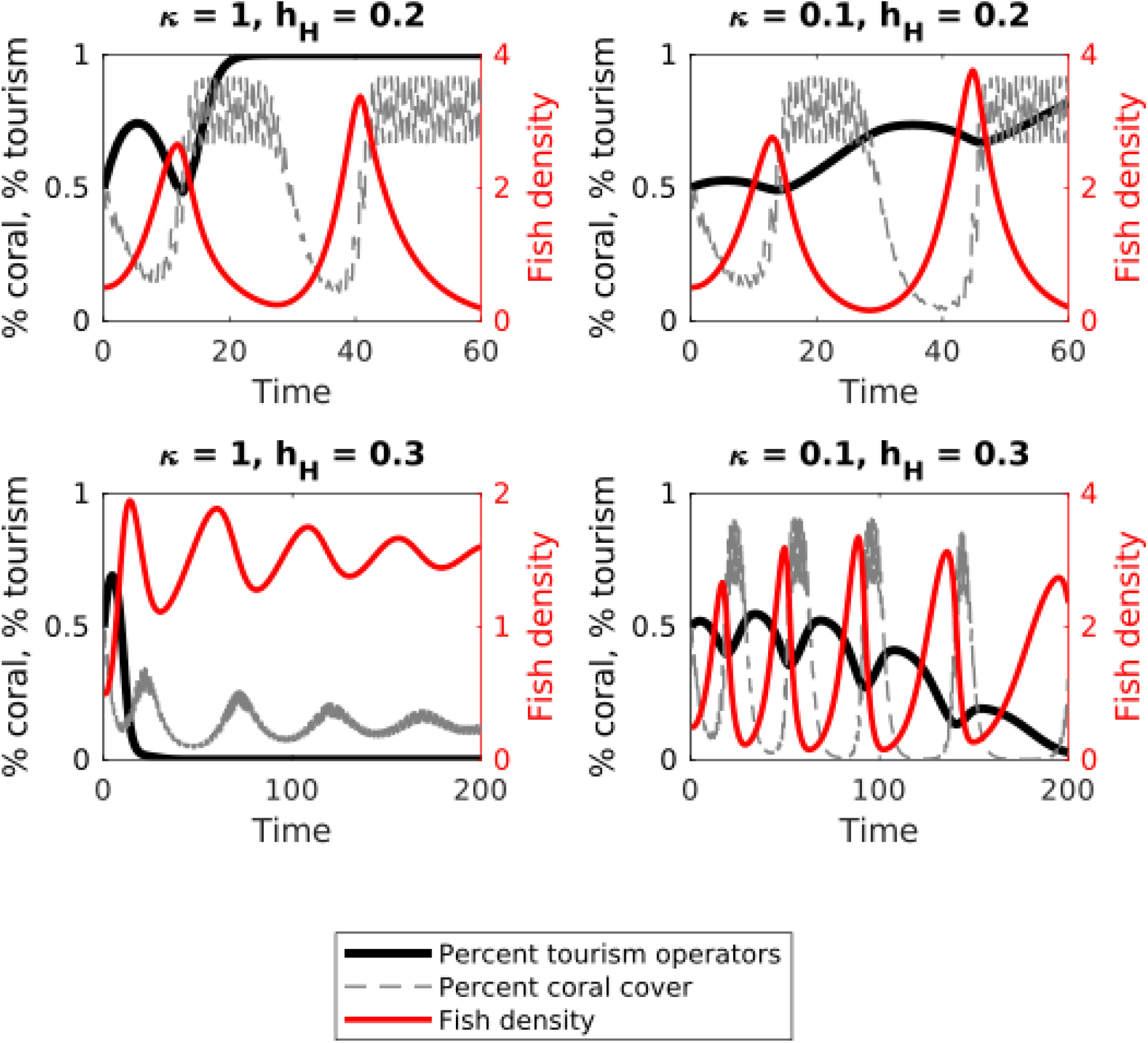
Time series showing coral cover, fish density and percentage of economic agents engaging in tourism for different values of *κ* and *h*_*H*_, within a single patch. The top two graphs show dynamics in the cyclic coral-dominant regime, while the bottom two show transient dynamics in the high-fish regime.

The boundaries between the different regimes are sharp, and transitions between the regimes can be driven by both overfishing and excessive nutrient loading (Fig. 2). Both the macroalgae-dominant and macroalgae-only regimes occur when economic activity converges to the fishing-only equilibrium, while the coral-dominant regime is coterminous with the area of parameter space in which economic activity converges to the tourism-only equilibrium. This indicates that in the macroalgae-dominant regime, some coral survived despite the fact that coral-related ecotourism was not economically viable.

### Spatial effects of local overfishing

We found that herbivorous fish and coral responded in opposite ways to the two patterns of local overfishing that we tested (Fig. 4). Herbivorous fish abundance was lower on average when overfished patches were contiguous (i.e. they were harmed more by habitat loss than fragmentation). In fact, the case where overfished patches alternated with non-overfished ones saw no decrease in average herbivorous fish abundance compared to the baseline. In contrast, coral had greater declines in the alternating-patch scenario, corresponding to habitat fragmentation as a result of overfishing. There, coral cover was uniformly low across the system, whereas in the contiguous-patch scenario large amounts of coral survived in the patches away from the stressed area.

**Figure 4:**
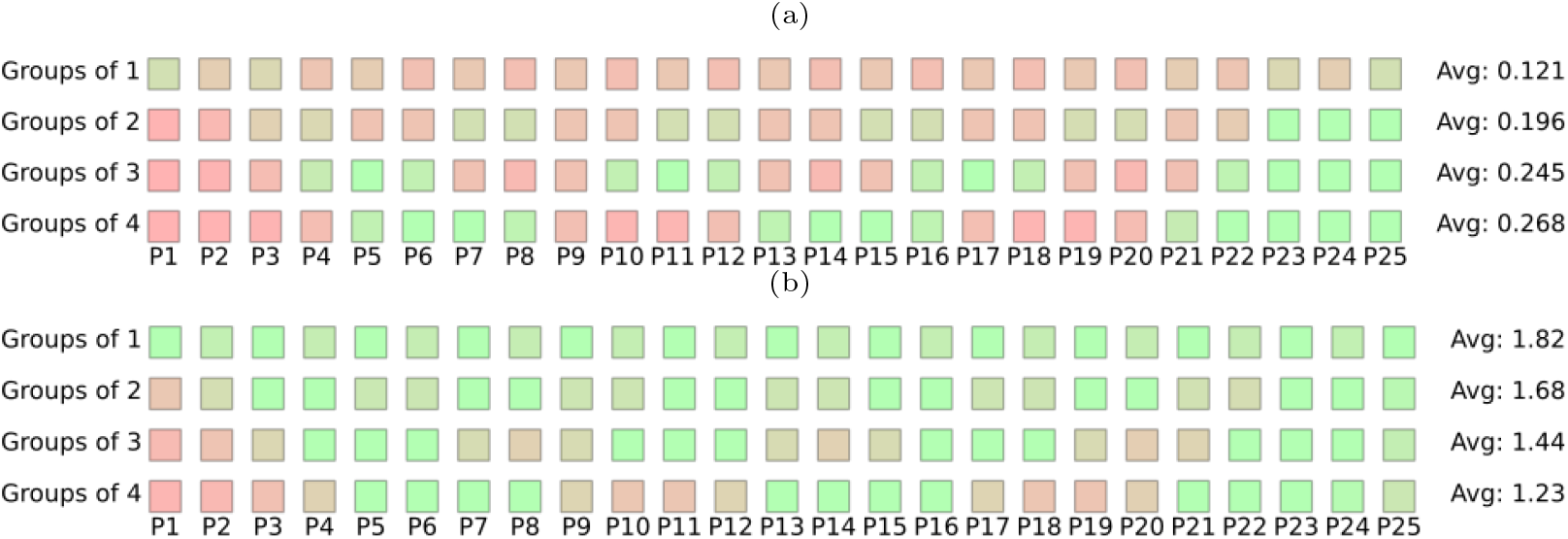
Coral cover (Fig. 4a) and fish density (Fig. 4b) in each of 25 patches, on a scale from red (low) to green (high), showing variation between contiguous and fragmented reefs. Here, 12 of 25 patches are overfished (arranged in groups of 1, 2, 3 and 4), and *σ*_*B*_ = 0.25. Averages of coral cover and fish density for the entire system are also provided for each configuration.

Increasing the proportion of patches that were overfished resulted in the expected linear decline in systemwide coral cover, as patches shifted one by one from being in the coral-dominant regime to the macroalgae-only regime. However, this masked nonlinear effects on coral in the overfished and non-overfished patches (Fig. 5). In the scenario where overfished patches were contiguous, coral cover fell off sharply in them but remained almost constant in the non-overfished ones. When overfished and non-overfished patches formed an alternating pattern, the decline in coral cover was steeper and was linear in both kinds of patches, and coral was completely extirpated at a ratio of two overfished patches for every one non-overfished one.

**Figure 5:**
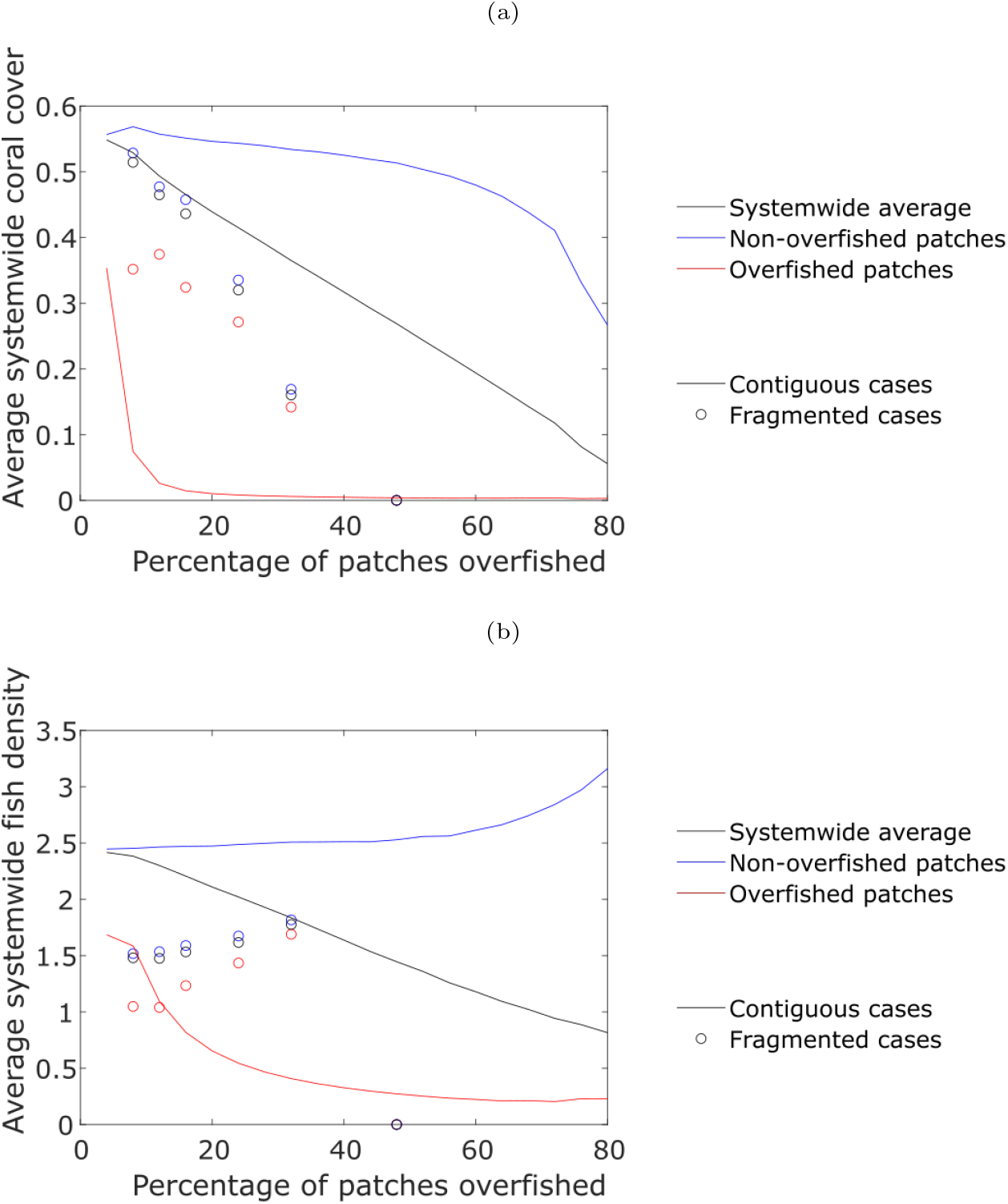
Average coral cover (Fig. 5a) and fish density (Fig. 5b) across a 25-patch system as a function of percentage of patches overfished, showing cases where overfished patches are contiguous and dispersed throughout the system. Values are shown for the system as a whole as well as for both overfished and non-overfished patches. Here, *σ*_*B*_ = 1.

As with habitat fragmentation, we found that the combination of nonlocal harvesting and the spillover effect had very different impacts on coral and herbivorous fish 7. For the case with contiguous strings of overfished and non-overfished patches, increasing fish dispersal ability (and hence the strength of any potential spillover effect) caused an increase of average fish density across the system by over 20 percent (Fig. 7b). This held regardless of how much time fishers spent locally. In the case where overfished and non-overfished patches alternated, and therefore any overfished patch could receive some spatial subsidies from an adjacent MPA, average fish density was higher than in the contiguous case (Fig. 4, Fig. 7b) with little dependence on fish dispersal ability in most cases. Coral dispersal ability had almost no effect on the abundance of fish or coral, with the exception of when both coral and fishing boats were almost entirely confined to their local patches. In contrast, we found that unauthorized fishing across MPA boundaries could lower coral cover by significant amounts, especially in the fragmented case (Fig. 7a) where we found losses of over thirty percent.

### Economic transitions

When we simulated systemwide transitions from a fishing-based to a tourism-based economy, we found that herbivorous fish returned to healthy levels after about 15 to 20 years, with the system returning to a coral-dominated state after an additional 10 years (Fig. 6). This was dependent on the degree to which coral was previously extirpated, as expected. Local economic transitions resulted in systemwide fish recovery when they occurred in as little as 12 percent of patches (Fig. 6b), and systemwide coral recovery happened when at least 56 percent of patches transitioned (Fig. 6a). (This meant that under some conditions, herbivorous fish were predicted to recover but coral was not.) The recovery times for fish and coral following these local transitions were typically longer by a few years than they were when the entire system transitioned, although the recovery times increased nonlinearly as the number of patches decreased towards the minimum number for which a recovery would take place, indicating the possibility of a bifurcation.

**Figure 6:**
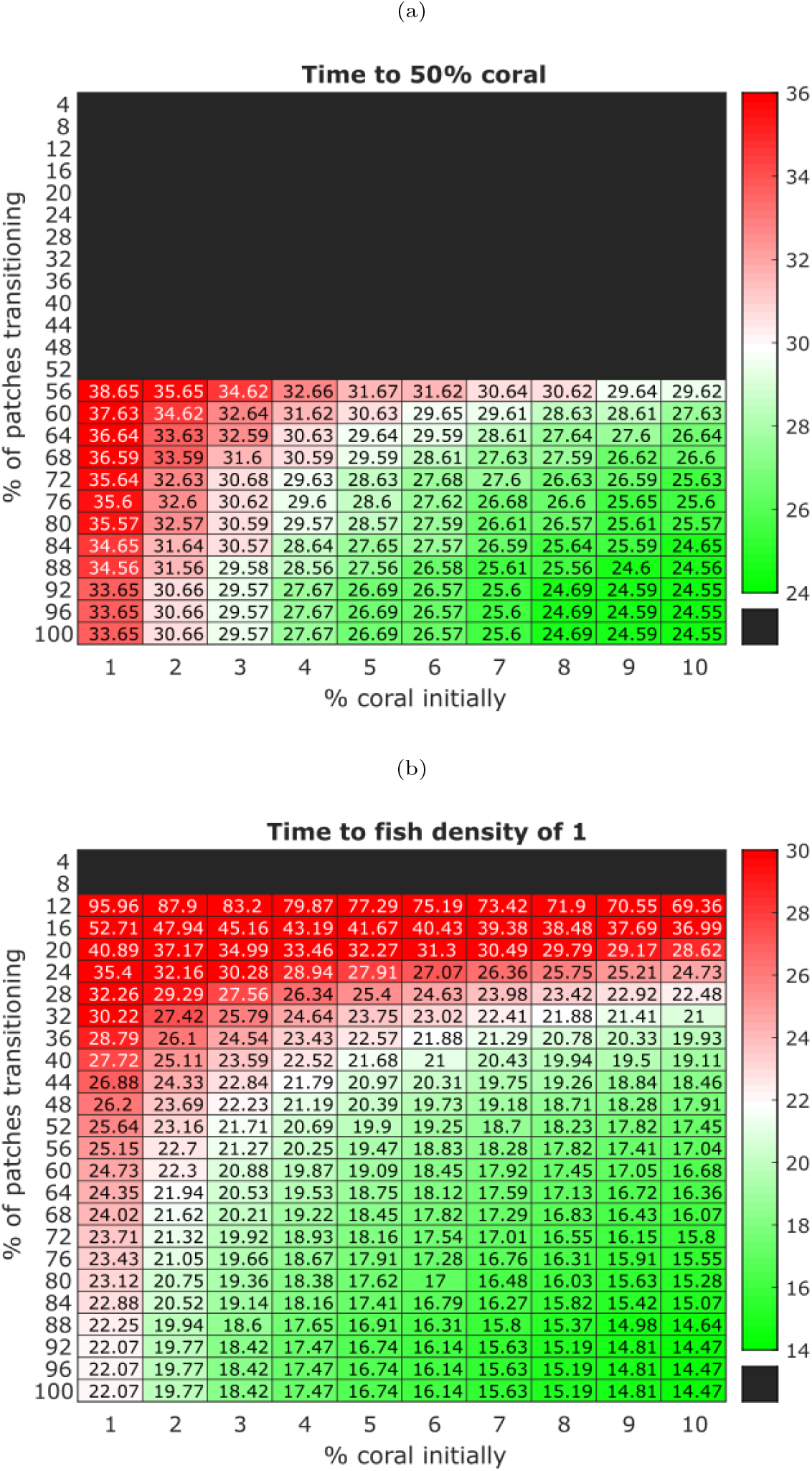
Time taken for average coral cover in a 25-patch system to return from a degraded state to 50 percent (Fig. 6a) and average fish density to return to 1 (Fig. 6b), following long-term economic transitions from fishing to tourism. Black boxes indicate that coral or fish was not observed to recover to the stated thresholds within 200 years.

**Figure 7:**
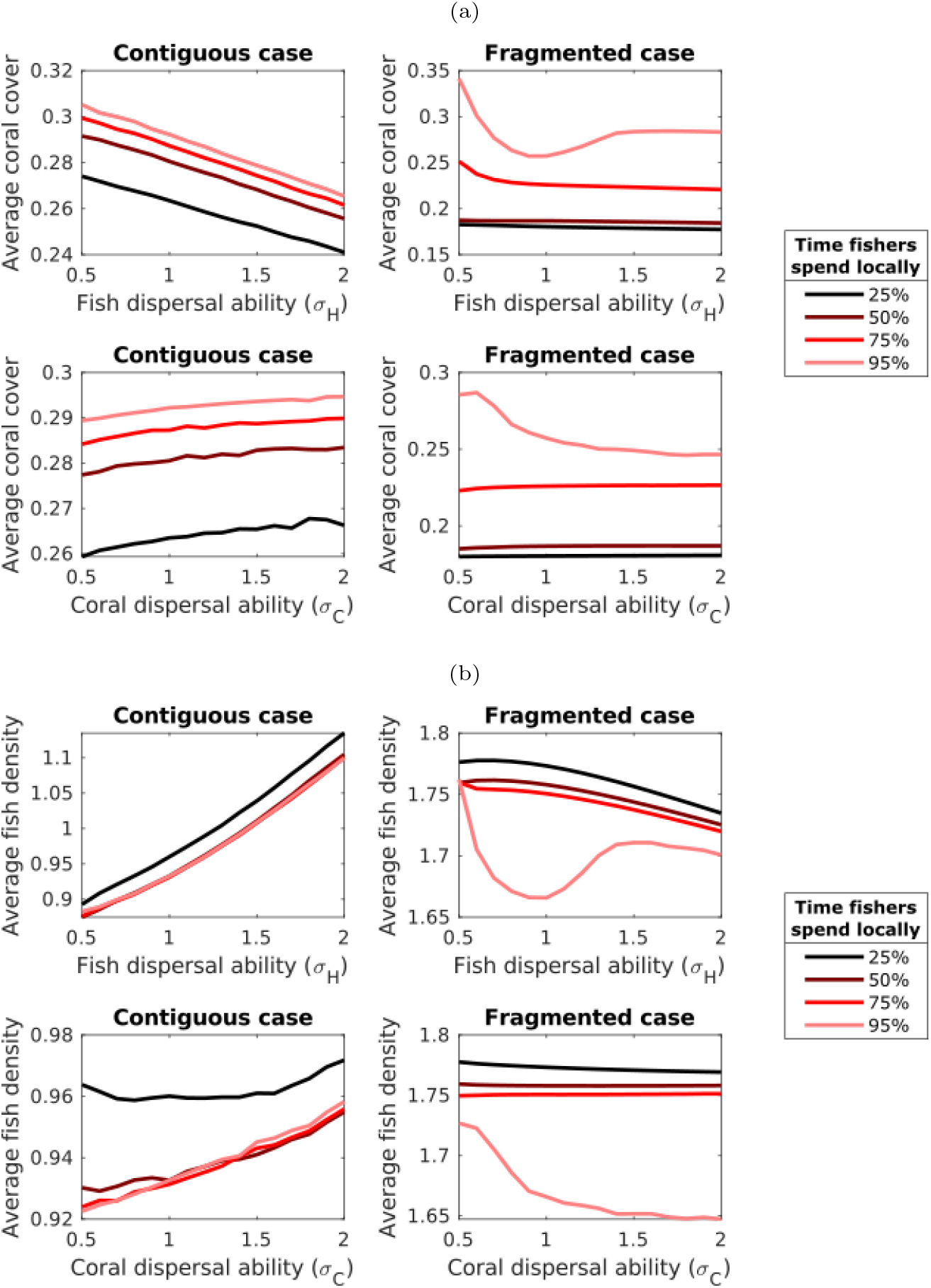
Average coral cover (Fig. 7a) and fish density (Fig. 7b) in a 25-patch system for different values of *σ*_*H*_, *σ*_*C*_ and *σ*_*B*_. Here, 13 of the 25 patches were overfished. Note the differences in scale on the vertical axes.

In our simulations involving short-term subsidization of the tourism sector in a heavily over-fished system, we found that depending on how much tourism was subsidized and for how long, four different outcomes were possible (Fig. 8). In increasing order of subsidy length or amount, these were the status quo (macroalgae dominance), a temporary recovery of the herbivorous fish population, a temporary recovery of both fish and coral, and a permanent shift to a tourism-based economy with healthy fish and coral populations. When fish and/or coral recovered, temporarily or permanently, this happened after the tourism subsidies had finished, indicating that tourism subsidies set off a positive feedback loop in terms of fish and coral populations.

**Figure 8:**
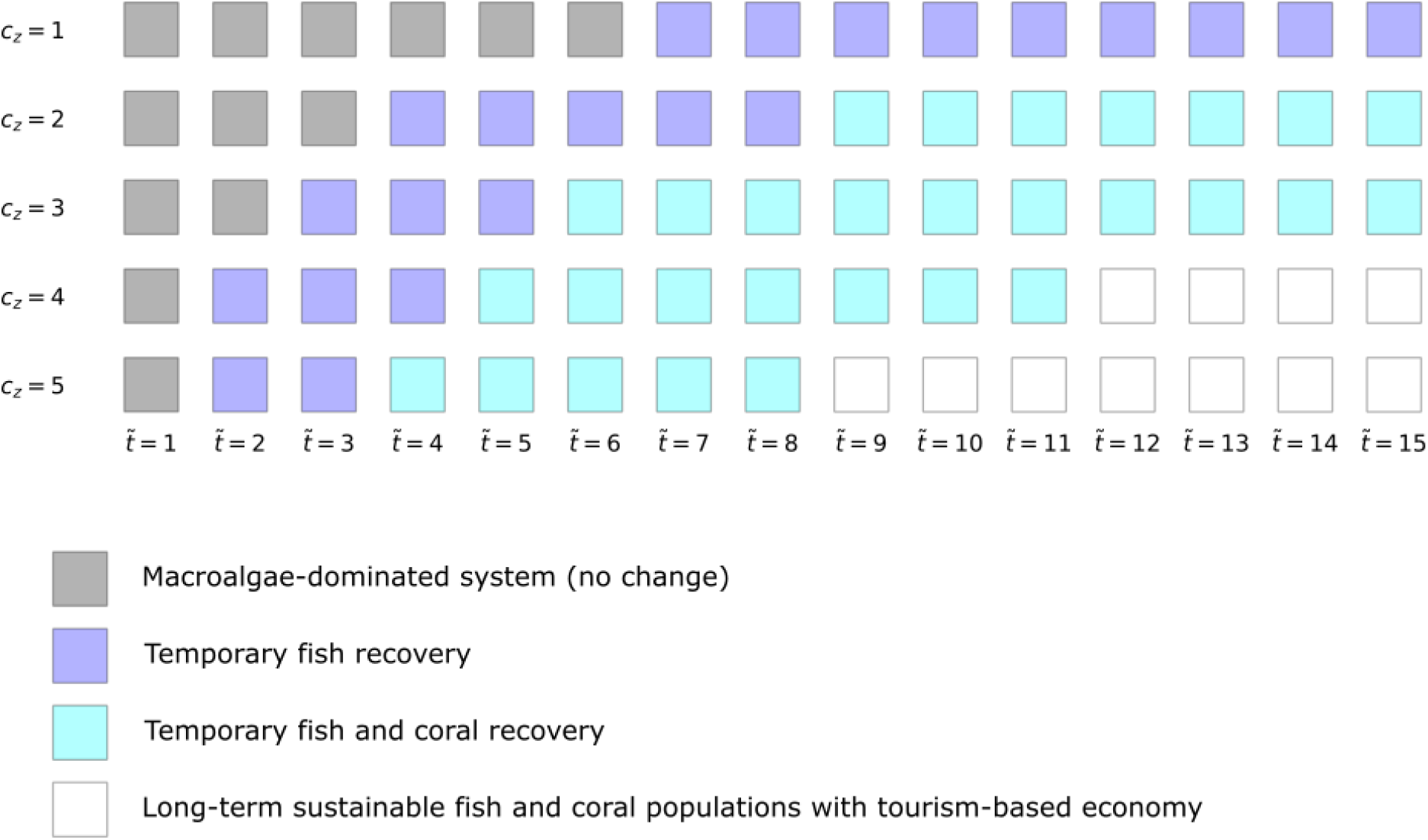
State reached by an overfished, macroalgae-dominated system after temporary subsidies to ecotourism. The four states shown differ in their transient and steady-state behaviour.

## Discussion

We found that habitat fragmentation (via overfishing and subsequent shift to conditions more favourable to macroalgae) strongly affected both coral and herbivorous fish. However, the effects of habitat fragmentation on these two functional groups were opposite to one another. When holding constant the percentage of patches that were overfished, coral cover was higher when long strings of non-overfished patches were adjacent to each other. In contrast, herbivorous fish populations were highest when overfished patches alternated with non-overfished ones, This is consistent with the results of Bonin et al. [10], who found that habitat fragmentation had only a temporary negative effect on reef fish, and when disentangled from habitat loss its long-term effects were neutral or even positive.

In a similar vein, we found that dispersal ability of herbivorous fish had much different effects on their abundance in contiguous and fragmented habitats. We found that the abundance of herbivorous fish was strongly dependent on their dispersal ability in scenarios where there were long stretches of overfished patches that could potentially receive spillover, although this saturates when MPAs and overfished patches alternate with each other and form a fragmented pattern. In these areas, herbivorous fish always exhibited a strong spillover effect, elevating the fish population size and hence the potential fish catch regardless of their dispersal ability (as there was always a place outside of any MPA for them to disperse into).

Our results suggest that a strategy of placing MPAs in the middle of overfished areas [68] would be effective in maximizing both fishing yield and standing fish populations, potentially by even more than the 20 percent increase predicted by Cabral et al. Our results also recommend the enforcement of MPA boundaries by requiring fishing operations to harvest mostly locally (above 75 percent in their home patches), as doing so is predicted to greatly boost coral populations while maintaining increased fish yield from the spillover effect. Although it has recently been shown that more mobile species show increased spillover tendencies and that some spillover is present regardless of whether habitats are fragmented or not [69], we believe that our study is the first to look at the interaction between these two factors.

The differing responses of coral and herbivorous fish to habitat fragmentation and dispersal ability can be explained by their different life history traits. When coral larvae disperse into an adjacent patch, if that patch is completely occupied by macroalgae, the coral larvae will be much less able to establish themselves. This can be seen in Fig. 7, where we found that coral larval dispersal ability has minimal effect on system coral cover. Our results here are in accordance with field results showing that the presence of macroalgae can inhibit both coral larval settlement and coral recruit survival after settlement [70, 71]. Hence, overfishing can cause coral to decline not just by removing predation pressure on faster-growing algae, but also by preventing colonization by coral larvae. This feedback loop can drive a shift to the macroalgae-dominant regime, and its presence explains why we found sharp regime boundaries (Fig. 2).

In the case when overfishing took place in a contiguous area, coral reacted very differently to an increase in the proportion of overfished patches depending on the local fishing rate. The average coral cover in overfished patches saw a steep dropoff after only a small number of patches became overfished, due to the breakdown of spatial subsidies. However, coral cover in the non-overfished patches (representing MPAs or areas fished at small-scale subsistence rates) remained at reasonably high levels, even when most patches were overfished and nonlocal harvesting was prevalent. This indicates the possibility of a conservation trap, in which a conservation-dependent species (in this case coral) is maintained via costly human intervention even though shifting the system to a more sustainable state would require less money and effort [72].

Given the predicted high discrepancy between coral health in overfished and non-overfished patches (or outside and inside MPAs) in the contiguous case, a manager could reasonably believe that only by implementing strict conservation measures can the coral be protected. However, we found two alternatives that may be considered if maintaining an MPA is not financially feasible. Firstly, the regime we found with high fish populations and stable coral cover (Fig. 2) features *z* converging to the fishing-only equilibrium, indicating that the coral population is not conservation-dependent. This is achievable for harvesting rates between 20 and 30 percent per year, similar to what has been seen in a previous model [50]. Secondly, we found that promoting ecotourism can shift a system back to a coral-dominated state over an appreciable timeframe, even if such promotion is temporary (Fig. 8) or spatially limited in scope (Fig. 6). These additional options allow coral reef managers more choice in the strategies they have for reef protection.

We found that following a large-scale economic transition that reduced fishing pressure on previously degraded reefs, fish could be expected to return to healthy levels after 14 to 20 years, with coral following about 10 years afterward. This is comparable to measured recovery times of reef ecosystems following other disturbances. For example, a recent long-term study on resilience of Caribbean coral and parrotfish populations found that percentage coral cover had risen from 36 to 47 percent, in line with pre-disturbance levels, seven years after a 2010 coral bleaching event [73]. Extending this rate of recovery of slightly less than two percent cover per year to the scenarios that we tested, which had much lower initial conditions for coral, yields recovery times very similar to what we found (Fig. 6). The same study found that parrotfish recovery after the disturbance, when assisted by a law enacted the same year that banned their harvesting, happened at a greater magnitude than coral recovery, echoing our findings that herbivorous fish recovery during an economic transition serves as a leading indicator for coral recovery.

Another long-term study found no significant increase in coral cover from low starting points (about 10 and 20 percent cover) in the six years following a period of disturbances [74], in accordance with our result that recovery from such levels should not be expected within that timeframe. Shifts to macroalgae-dominated regimes taking 14 years, about half the length we found for a shift in the other direction, have been observed in the field [18]; the difference can be explained by factors such as macroalgae’s ability to inhibit coral larval settlement (Fig. 7) and its higher intrinsic growth rate. Additionally, prior modelling results suggest that coral is able to recover after major hurricanes that happen once every 20 years, provided other environmental conditions are favourable [46, 75], which is also comparable to our results. This correspondence between our socially-driven transitions and the biologically-driven ones seen in previous field and modelling work helps validate the social component of our model, and indicates that coupled social-environmental interactions will be a useful addition to coral research going forward.

In addition to the spatial dynamics of coral and herbivorous fish, our results also show that different economic strategies (fishing and tourism) can coexist at a regional level. Specifically, we found that economic transitions from fishing to tourism along some parts of a reef can result in herbivorous fish populations rebounding across the system, enough so that fishing remains viable where the economic transitions did not take place. Our findings are supported by recent modelling results showing that fishing and tourism can coexist in the same area [76], as well as field observations that different economic strategies in marine communities have complex, overlapping distributions [77]. Similarly, our results suggest that reef health and fish catch can be effectively balanced by using strategies that we identified, such as selecting local areas in which tourism would be temporarily subsidized or annual harvesting rates would be limited to intermediate levels. As economic models of fishing that both take into account marine protected areas and are spatially explicit have only been put forward recently [78, 76], we believe that spatial modelling of economic strategies on coral reefs is an area ripe for future research.

## Tables

**Table 1:** Parameters, their units, and their associated values in this paper

| Parameter | Value | Units | Description |
| --- | --- | --- | --- |
| $r_H$ | 0.7 | $\text{yr}^{-1}$ | Herbivorous fish maximum intrinsic growth rate |
| $k_H$ | 0.5 | unitless | Half-saturation constant for herbivorous fish growth |
| $m_H$ | 0.1 | $\text{yr}^{-1}$ | Mortality rate for herbivorous fish from natural causes (i.e. non-harvesting) |
| $h_H$ | $0.05 - 0.5 - 0.8$ | $\text{yr}^{-1}$ | Commercial fish harvesting rate |
| $\tilde{h}_H$ | 0.05 | $\text{yr}^{-1}$ | Subsistence fish harvesting rate |
| $r_C$ | 5 | $\text{yr}^{-1}$ | Coral intrinsic growth rate |
| $m_C$ | 0.44 | $\text{yr}^{-1}$ | Coral mortality rate |
| $r_M$ | 12 | $\text{yr}^{-1}$ | Macroalgae intrinsic growth rate |
| $k_M$ | 80 | $\text{kmol N yr}^{-1}$ | Half-saturation constant for macroalgae growth |
| $m_M$ | 0.1 | $\text{yr}^{-1}$ | Macroalgae mortality rate |
| $\gamma$ | 1 | $\text{yr}^{-1}$ | Detritus decomposition rate |
| $q$ | $20 - 60 - 120$ | $\text{kmol N yr}^{-1}$ | Nitrogen loading rate |
| $\epsilon$ | 0.6 | $\text{yr}^{-1}$ | Nitrogen flushing rate |
| $f$ | 20 | $\text{kmol N}$ | Scaling constant for conversion of detritus into nutrients |
| $\kappa$ | 1 | $\text{yr}^{-1}$ | Rate at which economic agents can switch strategies |
| $c_z$ | $0 - 0 - 5$ | unitless | Economic utility for tourism (as compared to fishing) from external sources |
| $k_T$ | 1 | unitless | Scaling constant for how strongly tourism utility varies due to coral cover |
| $k_F$ | 1 | yr | Scaling constant for how strongly fishing utility varies due to fish catch |

## References

[1] D. R. Bellwood, T. P. Hughes, Regional-scale assembly rules and biodiversity of coral reefs, Science 292 (5521) (2001) 1532–1535. doi:10.1126/science.1058635.

[2] J. Veron, L. M. Devantier, E. Turak, A. L. Green, S. Kininmonth, M. Stafford-Smith, N. Peterson, Delineating the coral triangle, Galaxea, Journal of Coral Reef Studies 11 (2) (2009) 91–100. doi:10.3755/galaxea.11.91.

[3] M. L. Berumen, A. S. Hoey, W. H. Bass, J. Bouwmeester, D. Catania, J. E. M. Cochran, M. T. Khalil, S. Miyake, M. R. Mughal, J. L. Y. Spaet, P. Saenz-Agudelo, The status of coral reef ecology research in the red sea, Coral Reefs 32 (3) (2013) 737–748. doi:10.1007/s00338-013-1055-8.

[4] R. Costanza, R. d’Arge, R. de Groot, S. Farber, M. Grasso, B. Hannon, K. Limburg, S. Naeem, R. V. O’Neill, J. Paruelo, R. G. Raskin, P. Sutton, M. van den Belt, The value of the world’s ecosystem services and natural capital, Nature 387 (6630) (1997) 253–260. doi:10.1038/387253a0.

[5] C. M. Roberts, Effects of fishing on the ecosystem structure of coral reefs, Conservation Biology 9 (5) (1995) 988–995. doi:10.1046/j.1523-1739.1995.9051332.x-i1.

[6] J. W. McManus, L. A. Menez, K. N. Kesner-Reyes, S. G. Vergara, M. Ablan, Coral reef fishing and coral-algal phase shifts: implications for global reef status, ICES Journal of Marine Science 57 (3) (2000) 572–578. doi:10.1006/jmsc.2000.0720.

[7] T. R. McClanahan, C. C. Hicks, E. S. Darling, Malthusian overfishing and efforts to overcome it on kenyan coral reefs, Ecological Applications 18 (6) (2008) 1516–1529. doi:10.1890/07-0876.1.

[8] J. C. Blackwood, A. Hastings, P. J. Mumby, The effect of fishing on hysteresis in caribbean coral reefs, Theoretical Ecology 5 (1) (2010) 105–114. doi:10.1007/s12080-010-0102-0.

[9] M. J. Caley, K. A. Buckley, G. P. Jones, Separating ecological effects of habitat fragmentation, degradation and loss on coral commensals, Ecology 82 (12) (2001) 3435–3448. doi:10.1890/0012-9658(2001)082[3435:seeohf]2.0.co;2.

[10] M. C. Bonin, G. R. Almany, G. P. Jones, Contrasting effects of habitat loss and fragmentation on coral-associated reef fishes, Ecology 92 (7) (2011) 1503–1512. doi:10.1890/10-0627.1.

[11] L. A. Yeager, J. Estrada, K. Holt, S. R. Keyser, T. A. Oke, Are habitat fragmentation effects stronger in marine systems? a review and meta-analysis, Current Landscape Ecology Reports 5 (3) (2020) 58–67. doi:10.1007/s40823-020-00053-w.

[12] C. Mantyka, D. Bellwood, Macroalgal grazing selectivity among herbivorous coral reef fishes, Marine Ecology Progress Series 352 (2007) 177–185. doi:10.3354/meps07055.

[13] R. Ferrari, M. Gonzalez-Rivero, J. C. Ortiz, P. J. Mumby, Interaction of herbivory and seasonality on the dynamics of caribbean macroalgae, Coral Reefs 31 (3) (2012) 683–692. doi:10.1007/s00338-012-0889-9.

[14] M. O. Nadon, Stock assessment of the coral reef fishes of hawaii, 2016. (2017). doi:10.7289/V5/TM-PIFSC-60.

[15] R. Lennox, A. Filous, S. Cooke, A. Danylchuk, Substantial impacts of subsistence fishing on the population status of an endangered reef predator at a remote coral atoll, Endangered Species Research 38 (2019) 135–145. doi:10.3354/esr00942.

[16] J. C. Blackwood, A. Hastings, P. J. Mumby, A model-based approach to determine the long-term effects of multiple interacting stressors on coral reefs, Ecological Applications 21 (7) (2011) 2722–2733. doi:10.1890/10-2195.1.

[17] J. R. Zaneveld, D. E. Burkepile, A. A. Shantz, C. E. Pritchard, R. McMinds, J. P. Payet, R. Welsh, A. M. S. Correa, N. P. Lemoine, S. Rosales, C. Fuchs, J. A. Maynard, R. V. Thurber, Overfishing and nutrient pollution interact with temperature to disrupt coral reefs down to microbial scales, Nature Communications 7 (1) (jun 2016). doi:10.1038/ncomms11833.

[18] J. E. Arias-González, T. Fung, R. M. Seymour, J. R. Garza-Pérez, G. Acosta-González, Y.-M. Bozec, C. R. Johnson, A coral-algal phase shift in mesoamerica not driven by changes in herbivorous fish abundance, PLOS ONE 12 (4) (2017) e0174855. doi:10.1371/journal.pone.0174855.

[19] P. Dalzell, Catch rates, selectivity and yields of reef fishing, in: Reef Fisheries, Springer Netherlands, 1996, pp. 161–192. doi:10.1007/978-94-015-8779-2\_7.

[20] C. Kuster, V. Vuki, L. Zann, Long-term trends in subsistence fishing patterns and coral reef fisheries yield from a remote fijian island, Fisheries Research 76 (2) (2005) 221–228. doi:10.1016/j.fishres.2005.06.011.

[21] C. Birkeland, Symbiosis, fisheries and economic development on coral reefs, Trends in Ecology & Evolution 12 (9) (1997) 364–367. doi:10.1016/s0169-5347(97)01099-9.

[22] M. Fabinyi, The role of land tenure in livelihood transitions from fishing to tourism, Maritime Studies 19 (1) (2020) 29–39. doi:10.1007/s40152-019-00145-2.

[23] A. Diedrich, The impacts of tourism on coral reef conservation awareness and support in coastal communities in belize, Coral Reefs 26 (4) (2007) 985–996. doi:10.1007/s00338-007-0224-z.

[24] S. Grafeld, K. L. L. Oleson, L. Teneva, J. N. Kittinger, Follow that fish: Uncovering the hidden blue economy in coral reef fisheries, PLOS ONE 12 (8) (2017) e0182104. doi:10.1371/journal.pone.0182104.

[25] Q. T. K. Ngoc, Assessing the value of coral reefs in the face of climate change: The evidence from nha trang bay, vietnam, Ecosystem Services 35 (2019) 99–108. doi:10.1016/j.ecoser.2018.11.008.

[26] L. A. McKergow, I. P. Prosser, A. O. Hughes, J. Brodie, Regional scale nutrient modelling: exports to the great barrier reef world heritage area, Marine Pollution Bulletin 51 (1-4) (2005) 186–199. doi:10.1016/j.marpolbul.2004.11.030.

[27] K. ZŻ ychaluk, J. F. Bruno, D. Clancy, T. R. McClanahan, M. Spencer, Data-driven models for regional coral-reef dynamics, Ecology Letters 15 (2) (2011) 151–158. doi:10.1111/j.1461-0248.2011.01720.x.

[28] C. D. Storlazzi, M. van Ormondt, Y.-L. Chen, E. P. L. Elias, Modeling fine-scale coral larval dispersal and interisland connectivity to help designate mutually-supporting coral reef marine protected areas: Insights from maui nui, hawaii, Frontiers in Marine Science 4 (dec 2017). doi:10.3389/fmars.2017.00381.

[29] D. P. Thomson, R. C. Babcock, R. D. Evans, M. Feng, M. Moustaka, M. Orr, D. Slawinski, S. K. Wilson, A. S. Hoey, Coral larval recruitment in north-western australia predicted by regional and local conditions, Marine Environmental Research 168 (2021) 105318. doi:10.1016/j.marenvres.2021.105318.

[30] R. A. Abesamis, P. Saenz-Agudelo, M. L. Berumen, M. Bode, C. R. L. Jadloc, L. A. Solera, C. L. Villanoy, L. P. C. Bernardo, A. C. Alcala, G. R. Russ, Reef-fish larval dispersal patterns validate no-take marine reserve network connectivity that links human communities, Coral Reefs 36 (3) (2017) 791–801. doi:10.1007/s00338-017-1570-0.

[31] G. R. Almany, S. Planes, S. R. Thorrold, M. L. Berumen, M. Bode, P. Saenz-Agudelo, M. C. Bonin, A. J. Frisch, H. B. Harrison, V. Messmer, G. B. Nanninga, M. A. Priest, M. Srinivasan, T. Sinclair-Taylor, D. H. Williamson, G. P. Jones, Larval fish dispersal in a coral-reef seascape, Nature Ecology & Evolution 1 (6) (may 2017). doi:10.1038/s41559-017-0148.

[32] D. M. Beltrán, N. V. Schizas, R. S. Appeldoorn, C. Prada, Effective dispersal of caribbean reef fish is smaller than current spacing among marine protected areas, Scientific Reports 7 (1) (jul 2017). doi:10.1038/s41598-017-04849-5.

[33] J. C. Blackwood, C. Okasaki, A. Archer, E. W. Matt, E. Sherman, K. Montovan, Modeling alternative stable states in caribbean coral reefs, Natural Resource Modeling 31 (1) (2018) e12157. doi:10.1111/nrm.12157.

[34] T. Elmhirst, S. R. Connolly, T. P. Hughes, Connectivity, regime shifts and the resilience of coral reefs, Coral Reefs 28 (4) (2009) 949–957. doi:10.1007/s00338-009-0530-8.

[35] B. Spiecker, T. C. Gouhier, F. Guichard, Reciprocal feedbacks between spatial subsidies and reserve networks in coral reef meta-ecosystems, Ecological Applications 26 (1) (2016) 264–278. doi:10.1890/15-0478.

[36] C. Camargo, J. H. Maldonado, E. Alvarado, R. Moreno-Sánchez, S. Mendoza, N. Manrique, A. Mogollón, J. D. Osorio, A. Grajales, J. A. Sánchez, Community involvement in management for maintaining coral reef resilience and biodiversity in southern caribbean marine protected areas, Biodiversity and Conservation 18 (4) (2008) 935–956. doi:10.1007/s10531-008-9555-5.

[37] S. D. Jupiter, R. Weeks, A. P. Jenkins, D. P. Egli, A. Cakacaka, Effects of a single intensive harvest event on fish populations inside a customary marine closure, Coral Reefs 31 (2) (2012) 321–334. doi:10.1007/s00338-012-0888-x.

[38] D. M. P. Jacoby, F. Ferretti, R. Freeman, A. B. Carlisle, T. K. Chapple, D. J. Curnick, J. J. Dale, R. J. Schallert, D. Tickler, B. A. Block, Shark movement strategies influence poaching risk and can guide enforcement decisions in a large, remote marine protected area, Journal of Applied Ecology 57 (9) (2020) 1782–1792. doi:10.1111/1365-2664.13654.

[39] M. Fabinyi, The intensification of fishing and the rise of tourism: Competing coastal livelihoods in the calamianes islands, philippines, Human Ecology 38 (3) (2010) 415–427. doi:10.1007/s10745-010-9329-z.

[40] R. B. Cabral, S. D. Gaines, B. A. Johnson, T. W. Bell, C. White, Drivers of redistribution of fishing and non-fishing effort after the implementation of a marine protected area network, Ecological Applications 27 (2) (2017) 416–428. doi:10.1002/eap.1446.

[41] G. R. Almany, S. R. Connolly, D. D. Heath, J. D. Hogan, G. P. Jones, L. J. McCook, M. Mills, R. L. Pressey, D. H. Williamson, Connectivity, biodiversity conservation and the design of marine reserve networks for coral reefs, Coral Reefs 28 (2) (2009) 339–351. doi:10.1007/s00338-009-0484-x.

[42] L. W. Botsford, J. W. White, M.-A. Coffroth, C. B. Paris, S. Planes, T. L. Shearer, S. R. Thorrold, G. P. Jones, Connectivity and resilience of coral reef metapopulations in marine protected areas: matching empirical efforts to predictive needs, Coral Reefs 28 (2) (2009) 327–337. doi:10.1007/s00338-009-0466-z.

[43] B. J. Laurel, I. R. Bradbury, “big” concerns with high latitude marine protected areas (MPAs): trends in connectivity and MPA size, Canadian Journal of Fisheries and Aquatic Sciences 63 (12) (2006) 2603–2607. doi:10.1139/f06-151.

[44] A. C. Balbar, A. Metaxas, The current application of ecological connectivity in the design of marine protected areas, Global Ecology and Conservation 17 (2019) e00569. doi:10.1016/j.gecco.2019.e00569.

[45] M. J. Costello, D. W. Connor, Connectivity is generally not important for marine reserve planning, Trends in Ecology & Evolution 34 (8) (2019) 686–688. doi:10.1016/j.tree.2019.04.015.

[46] P. J. Mumby, A. Hastings, H. J. Edwards, Thresholds and the resilience of caribbean coral reefs, Nature 450 (7166) (2007) 98–101. doi:10.1038/nature06252.

[47] R. C. Babcock, J. M. Dambacher, E. B. Morello, É. E. Plagányi, K. R. Hayes, H. P. A. Sweatman, M. S. Pratchett, Assessing different causes of crown-of-thorns starfish outbreaks and appropriate responses for management on the great barrier reef, PLOS ONE 11 (12) (2016) e0169048. doi:10.1371/journal.pone.0169048.

[48] A. M. Szmant, Reproductive ecology of caribbean reef corals, Coral Reefs 5 (1) (1986) 43–53. doi:10.1007/bf00302170.

[49] T. Lindeberg, Scale-space for discrete signals, IEEE Transactions on Pattern Analysis and Machine Intelligence 12 (3) (1990) 234–254. doi:10.1109/34.49051.

[50] V. A. Thampi, M. Anand, C. T. Bauch, Socio-ecological dynamics of caribbean coral reef ecosystems and conservation opinion propagation, Scientific Reports 8 (1) (feb 2018). doi:10.1038/s41598-018-20341-0.

[51] R. Froese, D. Pauly, Fishbase, www.fishbase.org (2021).

[52] M. Abbiati, G. Buffoni, G. Caforio, G. D. Cola, G. Santangelo, Harvesting, predation and competition effects on a red coral population, Netherlands Journal of Sea Research 30 (1992) 219–228. doi:10.1016/0077-7579(92)90060-r.

[53] G. Santangelo, L. Bramanti, M. Iannelli, Population dynamics and conservation biology of the over-exploited mediterranean red coral, Journal of Theoretical Biology 244 (3) (2007) 416–423. doi:10.1016/j.jtbi.2006.08.027.

[54] A. Heyward, J. Collins, Growth and sexual reproduction in the scleractinian coral montipora digitata (dana), Marine and Freshwater Research 36 (3) (1985) 441. doi:10.1071/mf9850441.

[55] J. L. Ruesink, L. Collado-Vides, Modeling the increase and control of caulerpa taxifolia, an invasive marine macroalga, Biological Invasions 8 (2) (2006) 309–325. doi:10.1007/s10530-004-8060-3.

[56] B. E. Lapointe, M. M. Littler, D. S. Littler, A comparison of nutrient-limited productivity in macroalgae from a caribbean barrier reef and from a mangrove ecosystem, Aquatic Botany 28 (3-4) (1987) 243–255. doi:10.1016/0304-3770(87)90003-9.

[57] J. W. Fourqurean, J. C. Zieman, Nutrient content of the seagrass thalassia testudinum reveals regional patterns of relative availability of nitrogen and phosphorus in the florida keys usa, Biogeochemistry 61 (3) (2002) 229–245. doi:10.1023/a:1020293503405.

[58] B. E. Lapointe, R. A. Brewton, L. W. Herren, J. W. Porter, C. Hu, Nitrogen enrichment, altered stoichiometry, and coral reef decline at looe key, florida keys, USA: a 3-decade study, Marine Biology 166 (8) (jul 2019). doi:10.1007/s00227-019-3538-9.

[59] R. J. Lowe, J. L. Falter, Oceanic forcing of coral reefs, Annual Review of Marine Science 7 (1) (2015) 43–66. doi:10.1146/annurev-marine-010814-015834.

[60] C. E. Nelson, A. L. Alldredge, E. A. McCliment, L. A. Amaral-Zettler, C. A. Carlson, Depleted dissolved organic carbon and distinct bacterial communities in the water column of a rapid-flushing coral reef ecosystem, The ISME Journal 5 (8) (2011) 1374–1387. doi:10.1038/ismej.2011.12.

[61] P. M. Vitousek, J. D. Aber, R. W. Howarth, G. E. Likens, P. A. Matson, D. W. Schindler, W. H. Schlesinger, D. G. Tilman, Human alteration of the global nitrogen cycle: Sources and consequences, Ecological Applications 7 (3) (1997) 737–750. doi:10.1890/1051-0761(1997)007[0737:haotgn]2.0.co;2.

[62] M. F. Adame, M. E. Roberts, D. P. Hamilton, C. E. Ndehedehe, V. Reis, J. Lu, M. Griffiths, G. Curwen, M. Ronan, Tropical coastal wetlands ameliorate nitrogen export during floods, Frontiers in Marine Science 6 (nov 2019). doi:10.3389/fmars.2019.00671.

[63] S. Enríquez, C. M. Duarte, K. Sand-Jensen, Patterns in decomposition rates among photosynthetic organisms: the importance of detritus c:n:p content, Oecologia 94 (4) (1993) 457–471. doi:10.1007/bf00566960.

[64] S. Chidami, M. Amyot, Fish decomposition in boreal lakes and biogeochemical implications, Limnology and Oceanography 53 (5) (2008) 1988–1996. doi:10.4319/lo.2008.53.5.1988.

[65] M. Voss, H. W. Bange, J. W. Dippner, J. J. Middelburg, J. P. Montoya, B. Ward, The marine nitrogen cycle: recent discoveries, uncertainties and the potential relevance of climate change, Philosophical Transactions of the Royal Society B: Biological Sciences 368 (1621) (2013) 20130121. doi:10.1098/rstb.2013.0121.

[66] M. Pedersen, J. Borum, Nutrient control of estuarine macroalgae:growth strategy and the balance between nitrogen requirements and uptake, Marine Ecology Progress Series 161 (1997) 155–163. doi:10.3354/meps161155.

[67] B. Martínez, L. S. Pato, J. M. Rico, Nutrient uptake and growth responses of three intertidal macroalgae with perennial, opportunistic and summer-annual strategies, Aquatic Botany 96 (1) (2012) 14–22. doi:10.1016/j.aquabot.2011.09.004.

[68] R. B. Cabral, D. Bradley, J. Mayorga, W. Goodell, A. M. Friedlander, E. Sala, C. Costello, S. D. Gaines, A global network of marine protected areas for food, Proceedings of the National Academy of Sciences 117 (45) (2020) 28134–28139. doi:10.1073/pnas.2000174117.

[69] M. DiLorenzo, P. Guidetti, A. DiFranco, A. Calò, J. Claudet, Assessing spillover from marine protected areas and its drivers: A meta-analytical approach, Fish and Fisheries 21 (5) (2020) 906–915. doi:10.1111/faf.12469.

[70] I. Kuffner, L. Walters, M. Becerro, V. Paul, R. Ritson-Williams, K. Beach, Inhibition of coral recruitment by macroalgae and cyanobacteria, Marine Ecology Progress Series 323 (2006) 107–117. doi:10.3354/meps323107.

[71] F. J. Webster, R. C. Babcock, M. V. Keulen, N. R. Loneragan, Macroalgae inhibits larval settlement and increases recruit mortality at ningaloo reef, western australia, PLOS ONE 10 (4) (2015) e0124162. doi:10.1371/journal.pone.0124162.

[72] L. Cardador, L. Brotons, F. Mougeot, D. Giralt, G. Bota, M. Pomarol, B. Arroyo, Conservation traps and long-term species persistence in human-dominated systems, Conservation Letters 8 (6) (2015) 456–462. doi:10.1111/conl.12160.

[73] R. S. Steneck, S. N. Arnold, R. Boenish, R. de León, P. J. Mumby, D. B. Rasher, M. W. Wilson, Managing recovery resilience in coral reefs against climate-induced bleaching and hurricanes: A 15 year case study from bonaire, dutch caribbean, Frontiers in Marine Science 6 (jun 2019). doi:10.3389/fmars.2019.00265.

[74] P. Houk, D. Benavente, J. Iguel, S. Johnson, R. Okano, Coral reef disturbance and recovery dynamics differ across gradients of localized stressors in the mariana islands, PLoS ONE 9 (8) (2014) e105731. doi:10.1371/journal.pone.0105731.

[75] P. J. Mumby, A. Hastings, The impact of ecosystem connectivity on coral reef resilience, Journal of Applied Ecology 45 (3) (2007) 854–862. doi:10.1111/j.1365-2664.2008.01459.x.

[76] C. Falcó, H. V. Moeller, Optimal spatial management in a multiuse marine habitat: Balancing fisheries and tourism, Natural Resource Modeling (may 2021). doi:10.1111/nrm.12309.

[77] A. Ruiz-Frau, H. Hinz, G. Edwards-Jones, M. Kaiser, Spatially explicit economic assessment of cultural ecosystem services: Non-extractive recreational uses of the coastal environment related to marine biodiversity, Marine Policy 38 (2013) 90–98. doi:10.1016/j.marpol.2012.05.023.

[78] B. B. Xuan, C. W. Armstrong, Trading off tourism for fisheries, Environmental and Resource Economics 73 (2) (2018) 697–716. doi:10.1007/s10640-018-0281-5.

